# Succinate dehydrogenase activity supports de novo purine synthesis

**DOI:** 10.1101/2025.02.26.640389

**Authors:** Mushtaq A. Nengroo, Austin T. Klein, Heather S. Carr, Olivia Vidal-Cruchez, Umakant Sahu, Daniel J. McGrail, Nidhi Sahni, Peng Gao, John M. Asara, Hardik Shah, Marc L. Mendillo, Issam Ben-Sahra

## Abstract

The de novo purine synthesis pathway is fundamental for nucleic acid production and cellular energetics, yet the role of mitochondrial metabolism in modulating this process remains underexplored. In many cancers, metabolic reprogramming supports rapid proliferation and survival, but the specific contributions of the tricarboxylic acid (TCA) cycle enzymes to nucleotide biosynthesis are not fully understood. Here, we demonstrate that the TCA cycle enzyme succinate dehydrogenase (SDH) is essential for maintaining optimal de novo purine synthesis in normal and cancer cells. Genetic or pharmacological inhibition of SDH markedly attenuates purine synthesis, leading to a significant reduction in cell proliferation. Mechanistically, SDH inhibition causes an accumulation of succinate, which directly impairs the purine biosynthetic pathway. In response, cancer cells compensate by upregulating the purine salvage pathway, a metabolic adaptation that represents a potential therapeutic vulnerability. Notably, co-inhibition of SDH and the purine salvage pathway induces pronounced antiproliferative and antitumoral effects in preclinical models. These findings not only reveal a signaling role for mitochondrial succinate in regulating nucleotide metabolism but also provide a promising therapeutic strategy for targeting metabolic dependencies in cancer.

## Introduction

The tricarboxylic acid (TCA) cycle is a central metabolic hub that not only generates energy but also provides critical building blocks for biosynthetic processes^1^. Traditionally, the TCA cycle has been viewed primarily as an engine for oxidative phosphorylation, where the oxidation of nutrients produces reducing equivalents, such as NADH and FADH₂, that drive ATP production via the electron transport chain (ETC)^2^. This bioenergetic function depends on the continuous regeneration of electron acceptors; the ETC converts NADH back to NAD⁺ and FADH₂ back to FAD⁺, ensuring that the TCA cycle operates efficiently^1,3^.

However, beyond its role in energy production, the TCA cycle plays a vital role in anabolism^2,4^. It supplies key metabolites such as citrate and oxaloacetate, which serve as precursors for lipid and nucleic acid synthesis, respectively. In rapidly proliferating cells, the TCA cycle is critical for producing aspartate, which is a multifunctional amino acid that contributes to protein synthesis and acts as a major substrate for the de novo synthesis of both purines and pyrimidines^2,5^. This dual role highlights the importance of the TCA cycle in supporting the high biosynthetic demands of cancer cells^6–9^.

In recent years, an additional dimension of mitochondrial metabolism has emerged: the concept that TCA cycle intermediates also function as signaling molecules. These metabolites can modulate a variety of cellular processes, ranging from epigenetic regulation to stress responses. Despite their recognized importance, the mechanisms by which these metabolic by-products influence cellular pathways remain poorly defined^10–12^.

In this context, our study focuses on succinate, a key TCA cycle intermediate, and its role in nucleotide metabolism. We demonstrate that loss or inhibition of succinate dehydrogenase (SDH), a critical enzyme that links the TCA cycle to the ETC by oxidizing succinate to fumarate, leads to the accumulation of succinate. Remarkably, this buildup of succinate selectively impairs de novo purine synthesis while leaving pyrimidine synthesis largely intact. This selective inhibition suggests that succinate acts as a specific metabolic signal that disrupts the biosynthesis of purine nucleotides, a process essential for nucleic acid production and cell proliferation. The reduction in purine synthesis is accompanied by a compensatory upregulation of the purine salvage pathway, suggesting that cancer cells rely on this alternative route to maintain nucleotide pools when SDH is compromised. Furthermore, our preclinical data reveal that co-targeting SDH activity and the purine salvage pathway produces pronounced antiproliferative and antitumoral effects. By combining SDH inhibitors with agents that block the purine salvage pathway, we observed a synergistic reduction in cancer cell proliferation in vitro and tumor growth in vivo.

Overall, our research challenges the conventional view of mitochondria as mere energy producers, highlighting their critical role in orchestrating nucleotide biosynthesis and cellular proliferation. By showing that succinate accumulation disrupts de novo purine synthesis and drives compensatory purine salvage, our work lays the foundation for metabolism-targeted therapies that could improve outcomes in aggressive, treatment-resistant cancers.

## Results

### A coessentiality network analysis reveals links between succinate dehydrogenase complex and nucleotide metabolism

Genes with key roles in the same pathway are often most important for cellular fitness in the same contexts and thus can be identified as “coessential” when comparing gene dependencies across cancer cell lines^13^. To define the broader metabolic landscape regulated by succinate dehydrogenase (SDH) and its role in cancer cell fitness, we conducted a coessentiality network analysis using the FIREWORKS platform^14^. Focusing on the catalytic subunit of SDH, *SDHA*, we identified a striking coessentiality pattern between *SDHA* and several genes central to de novo nucleotide biosynthesis, including key enzymes involved in both purine (*MTHFD1*, *ATIC*, *PAICS*, *ADSL*, *GART*, *PFAS*) and pyrimidine (*CAD*, *DHODH*, *UMPS*) metabolism (Fig. 1A). This analysis highlights *SDHA* as a potential regulator of nucleotide synthesis, linking mitochondrial metabolism to the biosynthetic needs of proliferating cancer cells. Using DepMap’s CRISPR-Cas9 cell fitness (CERES) score data, we compared gene dependencies across a panel of more than 700 cancer cell lines. Loss of SDHA (*sgSDHA*) cells showed similar dependency patterns to the loss of other complex II subunits (*SDHB*, *SDHC*, *SDHD*) and, notably, to core purine biosynthesis genes (GART) and pyrimidine biosynthesis genes (*UMPS*), further supporting a functional interaction between SDH activity and nucleotide metabolism (Fig. 1B). To directly assess the impact of SDHA on cell proliferation, we generated SDHA knockout (SDHA KO) HeLa cells via CRISPR-Cas9 and validated SDHA loss by immunoblot (Fig. 1C). SDHA loss led to decreased proliferation of HeLa cells, which was fully rescued by the reintroduction of an SDHA cDNA construct, confirming that SDHA is required for optimal cell growth (Fig. 1C). We extended these findings to 786-O renal carcinoma cells, another model with high mitochondrial activity, and observed a significant reduction in cell proliferation upon SDHA knockout, indicating that SDHA’s role in supporting proliferation is not cell-line-specific (Fig. 1D). To generalize the decrease in cell fitness upon SDH inhibition, we treated a panel of cancer cell lines (H460, CAL-51, CT-26, and PC-3M) with the SDH inhibitor 3-nitropropionic acid (3-NPA), which suppresses the oxidation of succinate into fumarate^15^. Time-dependent reductions in cell proliferation were observed across all tested lines, with varying degrees of sensitivity, suggesting that some cancer cell types may be particularly reliant on intact SDH activity for growth (Fig. 1E, and Supplementary Fig. 1). To uncover pathways that may become essential upon SDH inhibition, we conducted a CRISPR metabolic screen using a focused metabolic phenotyping library targeting 232 metabolic genes covering all the major metabolic pathways (Supplementary Table 1). Cells were treated with vehicle or 3-NPA at the IC_20_ concentration for 21 days, followed by genomic DNA extraction and sequencing (Fig. 1F). Differential viability analysis revealed a striking increased dependence on purine synthesis genes including *PFAS* and *GMPS* in the presence of 3-NPA compared to vehicle control (Fig. 1G), further supporting a connection between SDH and nucleotide metabolism. Collectively, these findings revealed a previously unrecognized metabolic axis linking SDH activity to the de novo synthesis of both purines and pyrimidines, while functional assays demonstrated that *SDHA* loss or pharmacological inhibition leads to growth defects across diverse cancer models.

**Fig. 1.**
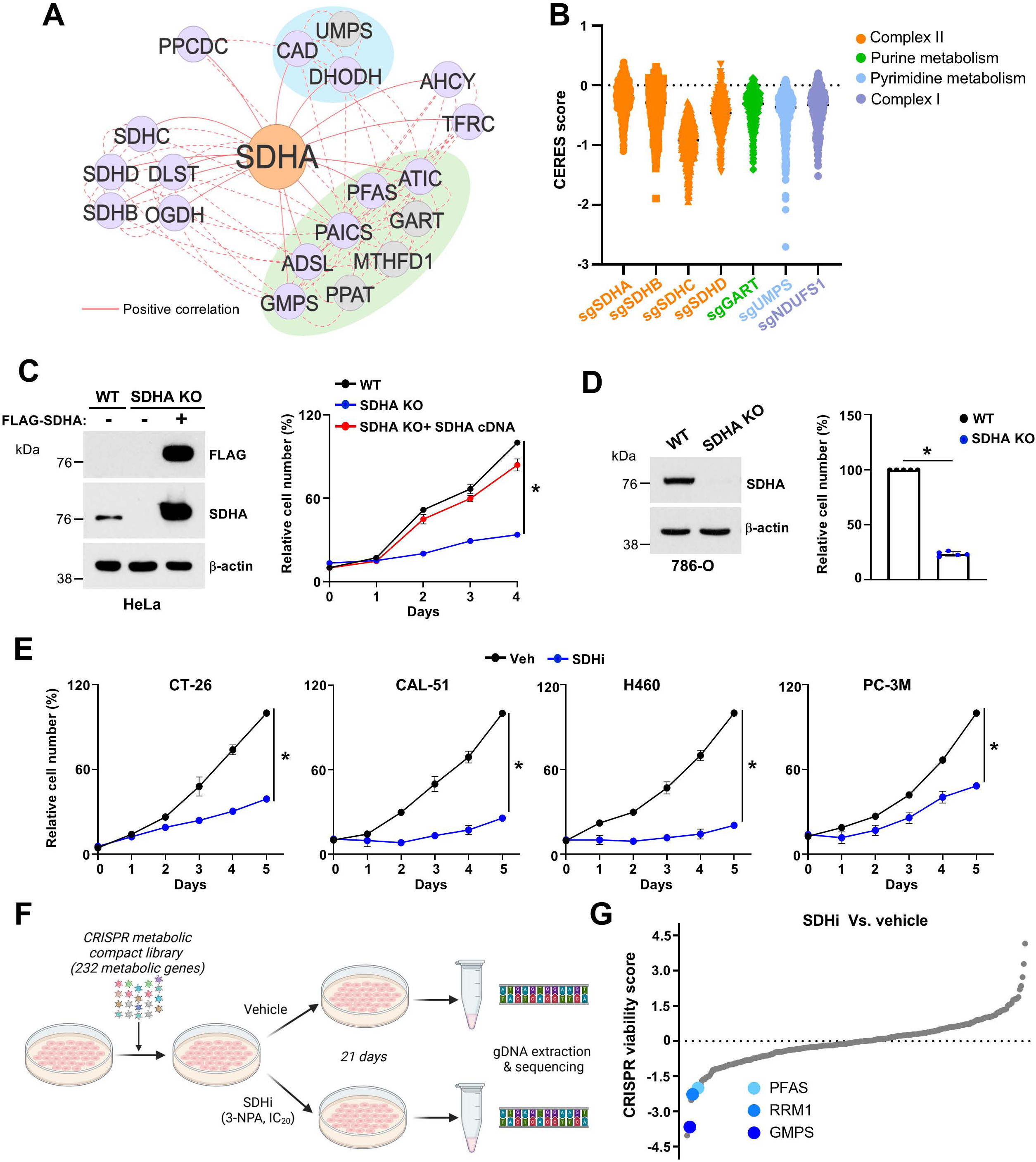
Coessentiality network and CRISPR-based screening identify SDH and purine metabolism as critical drivers of cancer cell proliferation. (A) a bottom-up coessentiality network analysis through the FIREWORKS portal between SDH subunits (SDHA, SDHB, SDHC, and SDHD) and other critical cellular metabolism genes. A coessentiality network of 20 positively correlated primary nodes with 5 positively correlated secondary nodes was generated. Solid red line represents a primary node that is positively coessential, and a dotted red line represents a secondary node that is positively coessential with a primary node. Note that double looping (two connections between a given gene pair) indicates that the correlation relationship is among the top-ranked for both genes at the specified rank thresholds (here, 20 for primary nodes and 5 for secondary nodes). The green module highlights genes involved in purine metabolism and the blue module highlights genes involved in pyrimidine metabolism. (B) CERES scores from DepMap for SDH subunits, GART (purine metabolism), UMPS (pyrimidine metabolism), and NDUFS1 (Complex I). (C) Immunoblot and relative viability of wildtype, SDHA knockout (SDHA KO), or SDHA KO HeLa cells reconstituted with a SDHA cDNA construct. (D) Immunoblot and relative cell viability of renal cancer cells wildtype or SDHA KO 786-O cells. (E) Time course treatment with 3-NPA (SDHi) in CT-26 (colorectal cancer (3NPA, 500 µM)), CAL-51 (breast cancer (3NPA, 200 µM)), H460 (lung cancer (3NPA, 200 µM)), and PC-3M (prostate cancer (3NPA, 500 µM)) cells. For all plots, data are shown as mean ± s.d. of n = 3 independent experiments. (F) Schematic of the CRISPR-based screen to identify metabolic genes required for growth under the LD_20_ concentration of 3-NPA. (G) Rank-order plot highlighting sgRNAs for purine genes in 3- NPA versus DMSO from the CRISPR–Cas9 screen. Data are representative of n = 3 independent biological replicates (C, D). \**P* < 0.05, by one-way ANOVA with Turkey’s post-hoc test for multiple comparisons (C) or unpaired t-test for pairwise comparisons (E, D).

### Succinate dehydrogenase activity supports de novo purine synthesis in vitro and in vivo

To define the role of SDH in nucleotide metabolism, we performed stable isotope tracing via liquid chromatography-mass spectrometry (LC-MS) approach. We employed two heavy isotope tracers [^15^N]-(amide)-glutamine and [^15^N-^13^C_2_]-glycine that are commonly used for studying the cellular activity of the de novo pyrimidine and purine synthesis pathways (Fig. 2A) ^16,17^. The incorporation of ^15^N into IMP (M+2) from [^15^N]-glutamine was significantly reduced in *SDHA*-deficient cells, while UMP (M+1) levels remained unchanged, suggesting that purine but not pyrimidine synthesis is compromised (Fig. 2B). Treatment with 3-NPA resulted in a similar reduction in the abundance of IMP (M+2), AMP (M+2), and GMP (M+3), confirming that SDH activity is necessary for de novo purine synthesis (Fig. 2C and Supplementary Fig. 2A, B). In contrast, pyrimidine intermediates, including N-carbamoyl-L-aspartate (M+1), orotate (M+1), and UMP (M+1), were not decreased upon SDH loss or inhibition (Fig. 2C, and Supplementary Fig. 2C, D). Moreover, aspartate incorporation into RNA, reflecting newly synthesized pyrimidines incorporated into RNA, was increased upon 3-NPA, while no changes was observed with the complex I inhibitor rotenone (Supplementary Fig. 1E), suggesting that SDH inhibition causes a differential effect on purine and pyrimidine metabolism. To more specifically track de novo purine synthesis activity, we measured the incorporation of [^15^N] and [^13^C] from [^15^N-^13^C_2_]-glycine into purine intermediates. While glycine uptake was not affected by SDH inhibition (Supplementary Fig. 2F), 3-NPA treatment reduced the abundance of AMP (M+3) and GMP (M+3), reinforcing the essential role of SDH in driving de novo purine synthesis (Fig. 2D). We next evaluated the impact of *SDHA* loss on nucleic acid synthesis derived from de novo purine synthesis by measuring the incorporation of U-[^14^C]-glycine into RNA. SDHA knockout or inhibition substantially decreased the incorporation of ^14^C into RNA in HeLa (Fig. 2E) and several other cell lines (Fig. 2F and Supplementary Fig. 2G, H), indicating reduced purine availability for nucleic acid synthesis. To extend these findings to an in vivo context, we established CT-26 or CAL-51 xenografts in mice and treated them with either vehicle or 3-NPA (Fig. 2G and Supplementary Fig. 2I). After tumors reached approximately 100 mm^3^, mice were administered [^13^C_2_-^15^N]-glycine, and tumors were harvested after 3 hours for LC-MS analysis. While glycine (M+3) abundance remained unaltered, SDH inhibition led to decreased abundance of IMP (M+3), AMP (M+3), and GMP (M+3) in colorectal and breast tumors (Fig. 2H and Supplementary Fig. 1J). These results confirm that SDH activity supports de novo purine biosynthesis in cells and tumors, highlighting the metabolic vulnerability induced by SDH inhibition.

**Fig. 2.**
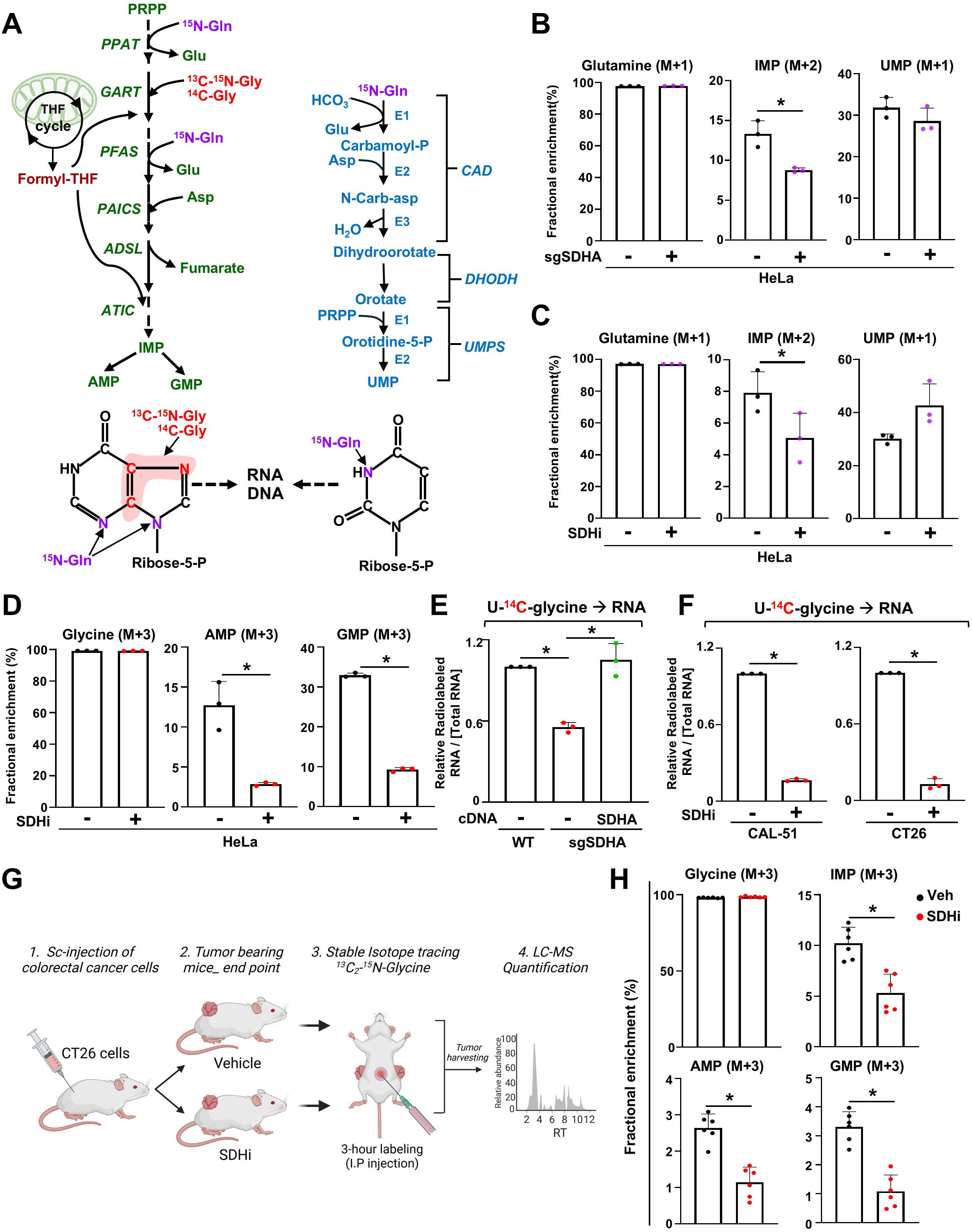
SDH activity is required to maintain optimal de novo purine nucleotide synthesis. (A) Schematic illustrating the enzymes and the different substrates required for the synthesis of purines and pyrimidines, which are used in the synthesis of nucleic acids. (B) Fractional enrichment (%) of ^15^N-labeled purine and pyrimidine intermediates, from wild-type or SDHA KO HeLa cells labeled with [^15^N-(amide)]-glutamine for 1 h. (C) Fractional enrichment (%) of ^15^N- labeled purine and pyrimidine intermediates from HeLa cells treated either with vehicle (DMSO) or 3-NPA (1 mM) and labeled with [^15^N-(amide)]-glutamine for 1 h. (D) Fractional enrichment (%) of ^15^N-^13^C-labeled purine intermediates, from vehicle or 3-NPA (1 mM) treated HeLa cells and labeled with [^15^N-^13^C_2_]–glycine for 4 h. (E) [U^14^C]-glycine radiolabeling was performed in wildtype and SDHA KO, and SDHA KO reconstituted with SDHA cDNA HeLa cells for 4 hours. RNA was extracted and radioactivity counts were measured. (F) Fractional enrichment (%) of labeled purine intermediates from vehicle (DMSO) or 3-NPA treated breast (CAL-51) and colorectal (CT-26) cancer cells, labeled with [^15^N-^13^C_2_]–glycine for 4 h. (G) Schematic illustrating the generation of CT26 xenograft tumors in the flanks of mice subsequently treated with vehicle or 3-NPA (30 mg/kg) prior to performing [^15^N-^13^C_2_]–glycine (0.5 g/kg) tracing in vivo. (H) Fractional enrichment (%) of ^15^N-^13^C-labeled glycine and purine intermediates from tumors are shown, n=5. For all plots, data are shown as mean ± s.d. of at least n = 3 independent biological replicates. \**P* <0.05. by two-tailed Student’s t-test (B-D, F, H) or one-way ANOVA with Tuckey’s post-hoc test for multiple pairwise comparisons (E).

### Aspartate synthesis does not underly the SDH-dependent regulation of purine synthesis

One of the essential anabolic molecules derived from the TCA cycle is aspartate, which serves as a key component for protein and nucleotide synthesis. Aspartate has mostly been primarily shown to be a major precursor for the synthesis of pyrimidine nucleotides because it provides 4 carbons and 1 nitrogen to the pyrimidine ring ^18,19^. Moreover, aspartate contributes to purine ring synthesis by providing an amino group, but no carbons.

To understand whether SDHA controls aspartate production, we performed [^13^C_6_]-glucose tracing experiments via LC-MS (Fig. 3A). While glucose uptake is increased in response to SDHA loss (Supplementary Fig. 3A), the abundance of aspartate (M+2) derived from glucose (M+6) was decreased upon 3-NPA treatment, suggesting that, as expected, SDH inhibition reduced aspartate synthesis (Fig. 3B). To determine whether a deficiency in mitochondrial aspartate synthesis, independently of SDH loss, can be sufficient to induce a decrease in de novo purine synthesis, we decided to inhibit mitochondrial aspartate synthesis by targeting enzymes downstream of SDH. In fact, aspartate is derived from oxaloacetate through a transamination reaction catalyzed by either the cytosolic glutamate-oxaloacetate transaminase (GOT1) enzyme or via GOT2 in the mitochondria^20,21^. As expected, loss of GOT2 (*sgGOT2*) slightly but significantly decreased the abundance of aspartate (M+4) derived from [^13^C_5_]-glutamine (Fig. 3D). Interestingly, GOT2 loss did not significantly impact purine synthesis (Fig. 3E), suggesting that a modest decrease in mitochondrial aspartate synthesis in *sgGOT2* cells is not sufficient to significantly alter de novo purine synthesis.

**Figure 3.**
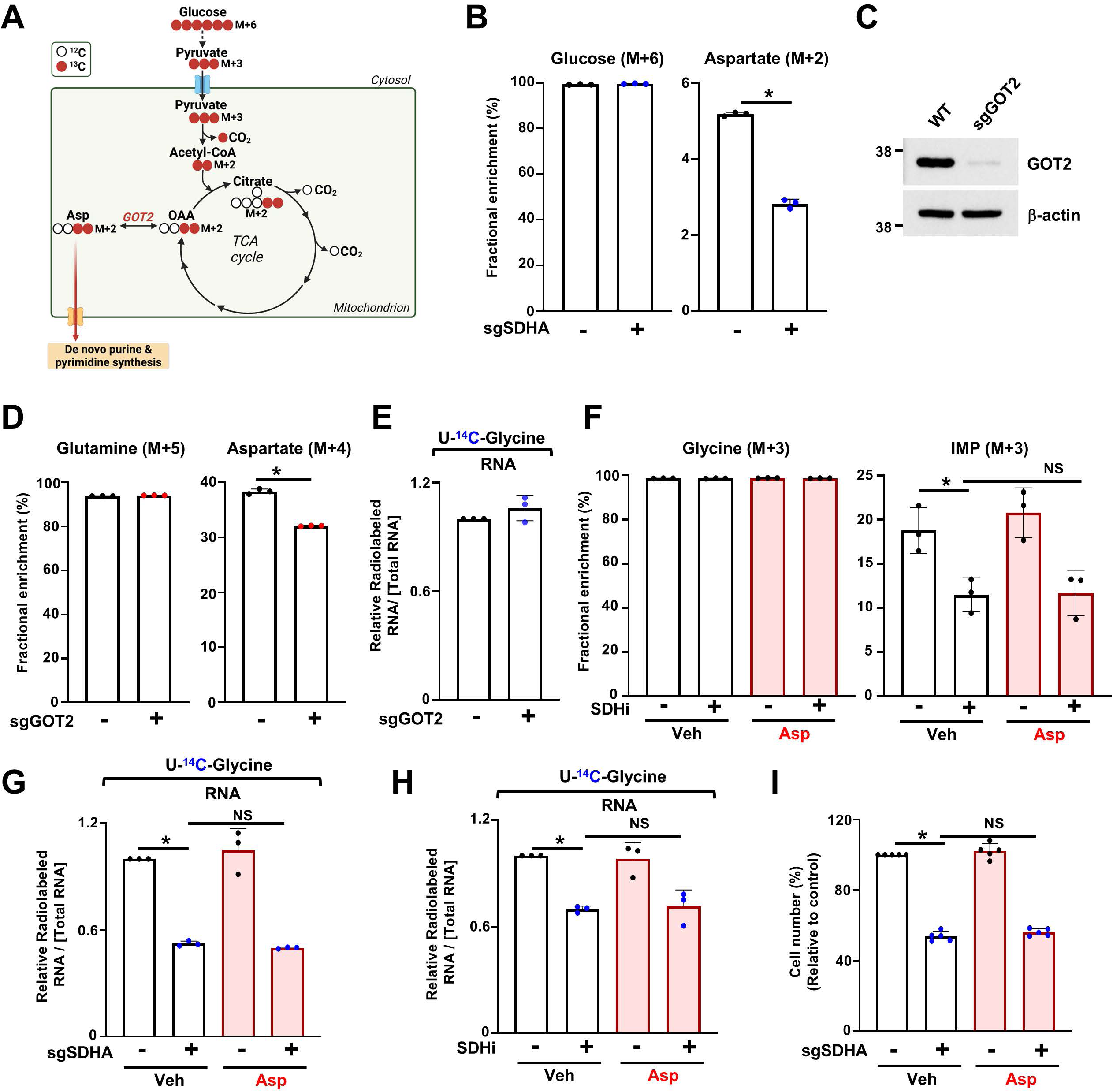
Depletion of aspartate upon SDH inhibition is not the cause for de novo purine synthesis inhibition. (A) Schematic illustrating the flow of carbons from [^13^C_6_]-glucose into the TCA cycle and aspartate synthesis. (B) Fractional enrichment (%) of the indicated isotopologues from wild-type and SDHA KO HeLa cells labeled with [^13^C_6_]-glucose for 2 h. Results show mean ± s.d., n=3 replicates per group. (C) Immunoblot of wildtype or GOT2 knockout (*sgGOT2*) HeLa cells. GOT2 and β-actin (loading control) are indicated. (D) Fractional enrichment (%) of the indicated isotopologues from wildtype and GOT2 KO HeLa cells labeled with [^13^C_5_]-glutamine (4 mM) for 2 h. (E) The relative levels of incorporation of [^14^C] into RNA are shown. Wild-type and GOT2 KO cells were labeled with [U-^14^C]-glycine for 4 hours, and the radioactivity measured reflects de novo purine synthesis. (F) Fractional enrichment (%) of indicated isotopologues from HeLa cells treated with either vehicle (DMSO) or SDHi (3-NPA) and labeled with [^15^N-^13^C_2_]– glycine (400 µM) in the presence or absence of aspartic acid (1 mM). (G) The relative levels of incorporation of [^14^C] into RNA are shown. Wildtype and SDHA KO cells were labeled for 4 hours in the presence or absence of aspartic acid (1mM). The radioactivity measured reflects de novo purine synthesis activity. (H) The relative levels of incorporation of [^14^C] into RNA are shown. HeLa cells were labeled for 6 hours and treated with either vehicle (DMSO) or 3-NPA in the presence or absence of aspartic acid (1 mM). (I) Wildtype and SDHA KO cells were cultured in the presence or absence of exogenous aspartate (1 mM) and cell number was measured via crystal violet staining after 96 h. For all plots, data are shown as mean ± s.d. of n = 3 independent biological replicates. \**P* < 0.05, by unpaired, two-tailed Student’s t-test (B, D) for pairwise comparisons or by one-way ANOVA coupled with Tukey’s post-hoc test for multiple comparison (F-I).

To measure the contribution of mitochondrial derived aspartate synthesis independently of glycolysis activity, we labeled cells with [^13^C_5_-]-glutamine in control cells or cells inhibited for SDH. Glutamine can be directly catabolized through the TCA cycle via the anaplerotic reaction that converts glutamate into α-ketoglutarate (Supplementary Fig. 3B). While SDH inhibition did not consistently alter the abundance of glutamate (M+5) derived from glutamine (M+5), the abundance of aspartate (M+4) was decreased, supporting that aspartate synthesis is inhibited in response to complex II loss or inhibition (Supplementary Fig. 3C). Moreover, SDH inhibition led to an increase in glutamine-dependent reductive carboxylation reflected by an increase in citrate (M+5) abundance, suggesting that complex II inhibition rewires the metabolic fate of glutamine towards anabolic precursor such as citrate (Supplementary Fig. 3B, C). To address whether the decrease in purine synthesis upon SDH inhibition was due to aspartate depletion, we performed [^15^N-^13^C_2_]-glycine labeling in cells supplemented or not with aspartate (1mM). We first confirmed that aspartate can be effectively transported in HeLa cells using [^3^H]-aspartate and that SDH loss or inhibition did not impede aspartate uptake (Supplementary Fig. 3D). We then observed that the abundance of IMP (M+3) was still decreased even when cells were exposed to exogenous aspartate (Fig. 3F). Similarly, exogenous aspartate did not rescue the inhibitory effects of SDH loss or inhibition on [^14^C] incorporation from glycine into RNA (Fig. 3G, H).

Similarly, the addition of aspartate did not rescue proliferation of cells inhibited for SDH (Fig. 3I). Overall, these results suggest that SDH activity is required to maintain mitochondrial aspartate production, but this mechanism did not contribute to the SDH-dependent regulation of de novo purine synthesis. Given that exogenous aspartate did not rescue purine synthesis and cell proliferation upon SDH inhibition, this indicates that there are other mechanisms that connect SDH activity to purine metabolism and growth of proliferating cells.

### Accumulation of succinate upon SDH inhibition leads to purine synthesis inhibition

To determine why, among other TCA cycle enzymes, SDH complex appears uniquely connected to the nucleotide metabolic network in our coessentiality analysis, we established knockout of each TCA cycle enzyme in HeLa cells using CRISPR-Cas9. We deleted citrate synthase (CS), aconitase 2 (ACO2), isocitrate dehydrogenase 3α (IDH3α), oxoglutarate dehydrogenase (OGDH), succinyl-CoA ligase (SUCLG1), fumarate hydratase (FH), and malate dehydrogenase 2 (MDH2) (Fig. 4A, B). We then assess how individual TCA enzyme disruptions impinge on de novo purine synthesis by quantifying the incorporation of [^14^C] from glycine into RNA. Among all knockouts, *SDHA* loss markedly reduced the incorporation of [^14^C] from glycine into RNA, signifying a specific role of SDH in supporting purine biosynthesis (Fig. 4C). SDHA loss caused a dramatic accumulation of succinate and a concurrent depletion of fumarate, and other TCA cycle intermediates, highlighting that SDH activity is essential for maintaining TCA cycle activity (Fig. 4D and Supplementary Fig. 4A). Nevertheless, supplementing SDH-deficient cells with cell-permeable fumarate (monomethylfumarate, MMF) did not restore purine synthesis, suggesting that diminished fumarate levels are not the root cause of the observed purine synthesis blockade (Supplementary Fig. 4B, C). Instead, succinate accumulation, which is rising from a baseline of 1–3 µM in wild-type cells to ∼1 mM upon SDH loss or inhibition, emerged as the significant culprit (Fig. 4E). To determine whether succinate accumulation is directly responsible for impaired purine synthesis, HeLa cells were treated with cell-permeable succinate (diethyl-succinate (DES)) (Supplementary Fig. 4D). Using ^13^C-DES, we confirmed that DES enters the cells and can be metabolized into succinate and fumarate (Supplementary Fig. 3E). Stable isotope tracing using [^15^N-^13^C_2_]-glycine revealed a marked decrease in labeled purine nucleotides, including AMP (M+3) and GMP (M+3), following DES treatment (Fig. 4F). DES treatment decreased the incorporation of [^14^C] from glycine into RNA, but did not suppress the incorporation of carbon and hydrogen from [C^3^H]-aspartate into RNA, further confirming that succinate accumulation impairs specifically the synthesis of purines but not that of pyrimidines (Fig. 4G and Supplementary Fig. 4F). These effects mimicked the metabolic consequences of SDH inhibition, strongly implicating elevated succinate as the inhibitory factor. These data establish that succinate accumulation, rather than fumarate depletion, is the primary cause of impaired de novo purine synthesis upon SDH inhibition. Elevated succinate acts as a metabolic inhibitor, reducing purine nucleotide production. These findings provide critical insight into how TCA cycle dysfunction can rewire nucleotide metabolism and highlight succinate as a key regulatory metabolite.

**Figure 4.**
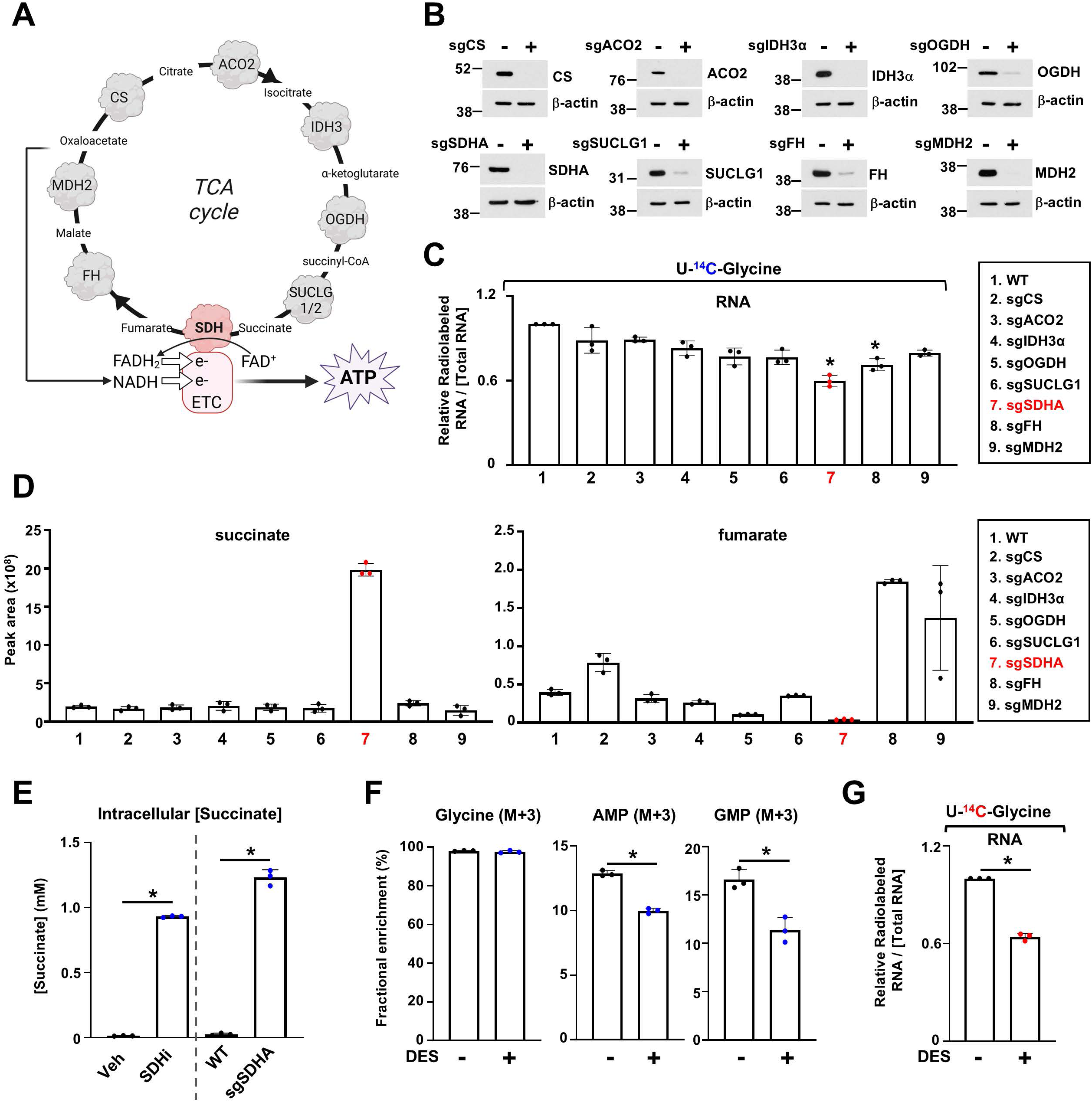
Accumulation of succinate upon SDH inhibition disrupts de novo purine synthesis. (A) Schematic of the TCA cycle and relevant enzymes. (B) Immunoblots of HeLa cells wildtype or knockout for CS (*sgCS*), ACO2 (*sgACO2*), IDH3α (*sgIDH3α*), OGDH (*sgOGDH*), SUCLG1 (*sgSUCLG1*), SDHA (*sgSDHA*), FH (*sgFH*), or MDH2 (*sgMDH2*). (C) The relative levels of incorporation of [^14^C] into RNA are shown. Wildtype or TCA enzyme KO cells were labeled with [U-^14^C]-glycine for 4 h, and radioactivity counts were measured, indicating de novo purine synthesis activity. (D) Normalized peak areas of succinate and fumarate in the indicated cells cultured in 10% dialyzed serum. (E) Absolute intracellular concentration of succinate after 8 h of 3-NPA (SDHi) treatment or in SDHA KO HeLa cells, compared with vehicle or wildtype conditions. (F) Fractional enrichment (%) of the indicated isotopologues from HeLa cells labeled with [^15^N- ^13^C_2_]–glycine for 4 h in the presence or absence of exogenous cell permeable succinate (diethyl- succinate (DES)). (G) The relative levels of incorporation of [^14^C] into RNA are shown. HeLa cells treated with either vehicle or DES (5 mM) for 8 h. For all plots, data are shown as mean ± s.d. of n = 3 independent biological replicates. \**P* < 0.05, by one-way ANOVA with Tukey’s post-hoc test (C), or unpaired, two-tailed Student’s t-test for pairwise comparisons (E-G).

### Co-targeting SDH activity and the purine salvage pathway impairs cancer cell proliferation and tumor growth

Building on our observation that succinate accumulation inhibits de novo purine synthesis and diminishes proliferation, we next explored whether cancer cells rely on the purine salvage pathway to overcome SDH loss. To test this possibility, we supplemented SDH-inhibited cells with exogenous purine nucleobases and nucleosides (hypoxanthine and inosine) and found that this addition fully restored cell proliferation (Fig. 5A). This result demonstrated that the growth defect caused by SDH inhibition can be rescued by providing alternative purine precursors. To confirm that the purine salvage pathway underpins this rescue, we performed stable isotope tracing using [^13^C_5_]-hypoxanthine and measured the incorporation of labeled carbon into purine intermediates (Fig. 5B). In SDH-deficient cells, the uptake of hypoxanthine and its incorporation into downstream purine intermediates such as AMP and GMP were significantly increased (Fig. 5C), highlighting a shift toward the purine salvage pathway as a compensatory mechanism. Further validation using [^3^H]-hypoxanthine incorporation assays showed enhanced labeling of RNA in SDH-deficient cells (Fig. 5D), which was reversed upon re-expression of wild-type SDHA, confirming that the observed effect was directly linked to SDH loss. Given that SDH inhibition promotes reliance on the purine salvage pathway, we tested whether co-targeting SDH and the purine salvage pathway would more effectively suppress cancer cell proliferation. We treated H460 (lung cancer), CT-26 (colorectal cancer), HeLa (cervical cancer), and PC-3M (prostate cancer) cells with low doses of 3-NPA and the purine salvage pathway inhibitor 6-mercaptopurine (6-MP), either alone or in combination. While single-agent treatments had modest effects, the combination of 3-NPA and 6-MP substantially curtailed cancer cell proliferation (Fig. 5E, F, and Supplementary Fig. 5A, B). A similar reduction was observed when cells were treated with cell-permeable succinate (DES) in combination with 6-MP (Fig. 5G). To translate these in vitro findings into a physiologically relevant system, we employed an immunocompetent mouse model of colorectal cancer using CT-26 cells. This syngeneic BALB/c model closely mirrors clinical tumor-immune interactions^22^. Mice were injected subcutaneously with CT-26 cells, and once tumors were established, they were randomized into four treatment groups: vehicle, 3-NPA, 6-MP, or the combination of 3-NPA and 6-MP. Tumor growth was monitored over 21 days, and mouse health was assessed by tracking body weight as a general indicator of toxicity (Fig. 5H).

**Figure 5.**
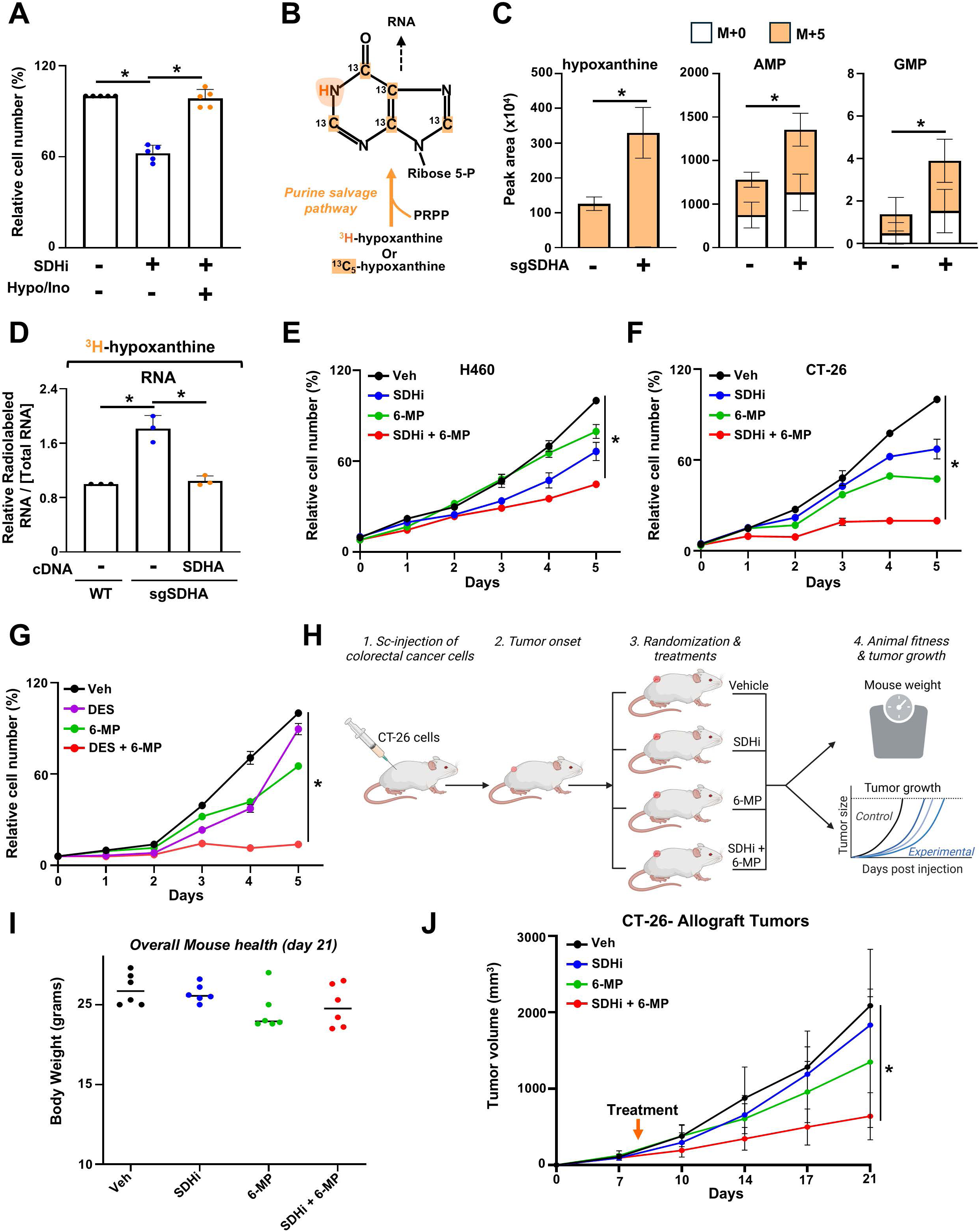
Co-targeting SDH and purine salvage creates a metabolic vulnerability for tumor growth. (A) CAL-51 cells treated with 3-NPA (100 µM), with or without exogenous hypoxanthine (100 µM) or inosine (100 µM). Crystal violet assay was performed after 72 h. (B) Schematic of the purine salvage pathway, highlighting ^13^C or ^3^H incorporation from hypoxanthine into IMP and subsequently into RNA. (C) Normalized peak areas of the indicated metabolites, measured via LC-MS, from HeLa cells labeled with ^13^C_5_-hypoxanthine for 3 hours. (D) Incorporation of [^3^H] from [^3^H]-hypoxanthine into RNA is shown. Labeling in wildtype, SDHA KO, or SDHA KO reconstituted with SDHA cDNA HeLa cells was performed for 4 h. Radioactivity counts were measured. (E) H460 cells treated with either vehicle, 3-NPA (30 µM), 6-MP (12.5 µM), or the combination, and crystal violet assays were performed every 24 h. (F) CT-26 cells treated with either vehicle, 3- NPA (250 µM), 6-MP (2 µM), or the combination, and crystal violet assays were performed every 24 h. (G) HeLa cells treated with vehicle or diethyl-succinate (DES, 3 mM), 6-MP (3 µM), or the combination, and crystal violet assays were performed every 24 h. (H) Schematic demonstrating the generation of CT-26 subcutaneous allograft tumors in Balb/c mice and the treatment regimen. (I) Mouse body weight at study endpoint, reflecting overall health. (J) Growth curves of CT-26 xenograft tumors treated with vehicle, 3-NPA (30 mg/kg), 6-MP (15 mg/kg), or their combination (n = 6). Data represent mean ± s.d. of at least n = 3 (A, C-F) or n =6 (H, I) biological replicates. * *P* < 0.05, by unpaired, two-tailed Student’s t-test (C) or one-way ANOVA with Tukey’s post-hoc test (A, D-G, J).

While single-agent treatments modestly affected tumor progression, the combination of 3-NPA and 6-MP significantly suppressed tumor growth without causing notable toxicity, as indicated by stable body weights (Fig. 5I, J). Overall, these findings reveal that while SDH inhibition reduces de novo purine synthesis, cancer cells compensate by upregulating the purine salvage pathway to maintain nucleotide pools and sustain proliferation. However, co-targeting SDH activity and the purine salvage pathway creates a synthetic lethal interaction that effectively impairs tumor growth, offering a promising therapeutic approach. This combinatorial strategy exploits a metabolic vulnerability in cancer cells and supports the rationale for developing dual-targeting therapies to improve outcomes in tumors with dysregulated mitochondrial metabolism.

## Discussion

We find a metabolic connection between mitochondrial complex II and de novo purine synthesis, revealing that SDH activity is critical not only for bioenergetics but also for sustaining the anabolic demands of rapidly proliferating cancer cells. We demonstrate that genetic deletion or pharmacological inhibition of SDH leads to a pronounced accumulation of succinate, which in turn selectively impairs de novo purine synthesis while sparing pyrimidine production. This specificity indicates that succinate acts as a targeted metabolic signal, directly inhibiting key enzymes involved in purine biosynthesis. Although steady-state purine nucleotide levels may appear unchanged, dynamic isotope tracing reveals a rapid, succinate-induced blockade of de novo purine synthesis, highlighting the transient yet critical nature of this metabolic regulation that ultimately results in reduced RNA synthesis and impaired cell proliferation.

Furthermore, the metabolic stress imposed by SDH inhibition triggers a compensatory upregulation of the purine salvage pathway, owing to the cells’ need to restore nucleotide pools and maintain growth despite compromised de novo synthesis. Previous metabolic analyses have shown that both de novo and salvage pathways comparably contribute to purine nucleotide pools in tumors^23^, emphasizing how reliance on salvage increases when de novo synthesis is blocked by succinate accumulation. The synthetic lethal interaction achieved by co-targeting SDH activity and the purine salvage pathway suggests that cancer cells have limited capacity to compensate for dual metabolic disruption. Our preclinical data, obtained from both in vitro experiments and in vivo xenograft models, support the therapeutic potential of this combinatorial approach, as it significantly reduces tumor growth without causing overt toxicity.

These findings challenge the conventional view of mitochondria solely as energy producers and emphasize their pivotal role in regulating biosynthetic pathways through metabolite signaling^24^. The dual role of succinate, both as an intermediary in the TCA cycle and as a signaling molecule that can modulate nucleotide synthesis, provides an original framework for understanding cancer cell metabolism. In particular, our study reveals that mitochondrial dysfunction, through SDH inhibition, can lead to a metabolic vulnerability that is therapeutically exploitable. Our work extends previous observations by Wu et al., owing to their demonstration that complex I impairment shifts lung cancer cells toward purine salvage, and aligns with recent findings that fumarate accumulation in FH-deficient tumors similarly blocks de novo purine synthesis^25,26^. Together, these studies suggest a convergent metabolic vulnerability across distinct mitochondrial defects. Beyond its impact on nucleotide synthesis, succinate accumulation is known to influence other cellular processes, such as epigenetic reprogramming via TET demethylase inhibition and hypoxia-inducible factor (HIF) stabilization, owing to its ability to interfere with key regulatory enzymes^27,28^. These broader effects may contribute to human diseases, inflammation, and therapy resistance, highlighting the multifaceted role of mitochondrial metabolites in biology ^29–31^.

In summary, our research uncovers a previously underappreciated regulatory axis linking SDH activity to nucleotide metabolism. By demonstrating that succinate accumulation disrupts de novo purine synthesis and forces a reliance on the purine salvage pathway, we provide a compelling rationale for targeting these metabolic processes in cancer therapy. Future work will focus on refining dual-inhibition strategies, exploring additional compensatory pathways, and validating these findings in clinical contexts. This study not only advances our understanding of mitochondrial metabolism in cancer but also opens avenues for the development of targeted, metabolism-based therapies for aggressive, treatment-resistant tumors.

## Supplementary Figures

**Supplementary Fig. 1.**
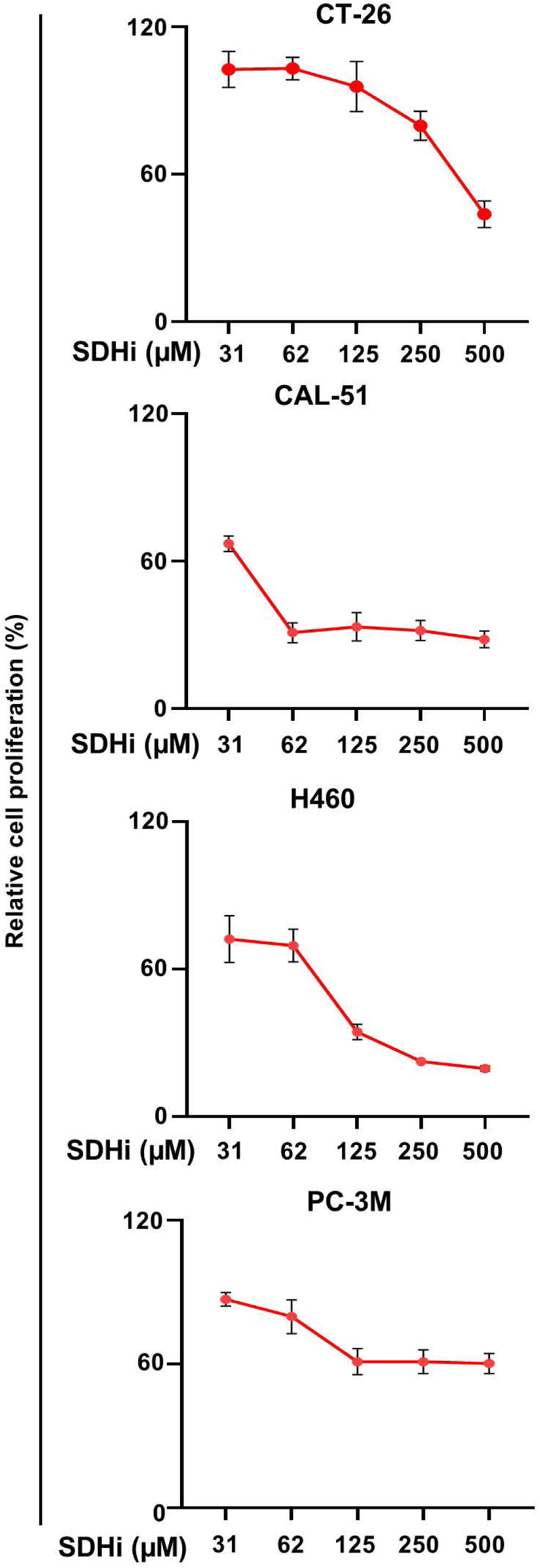
Dose-dependent SDH inhibition by 3-NPA reduces proliferation in multiple cancer cell lines. Shown are dose–response curves for 3-NPA (SDHi) treatment in CT- 26, CAL-51, H460, and PC-3M cells, illustrating decreased cell proliferation with increasing inhibitor concentrations. Data represent mean ± s.d. from n = 3 independent biological replicates.

**Supplementary Fig. 2.**
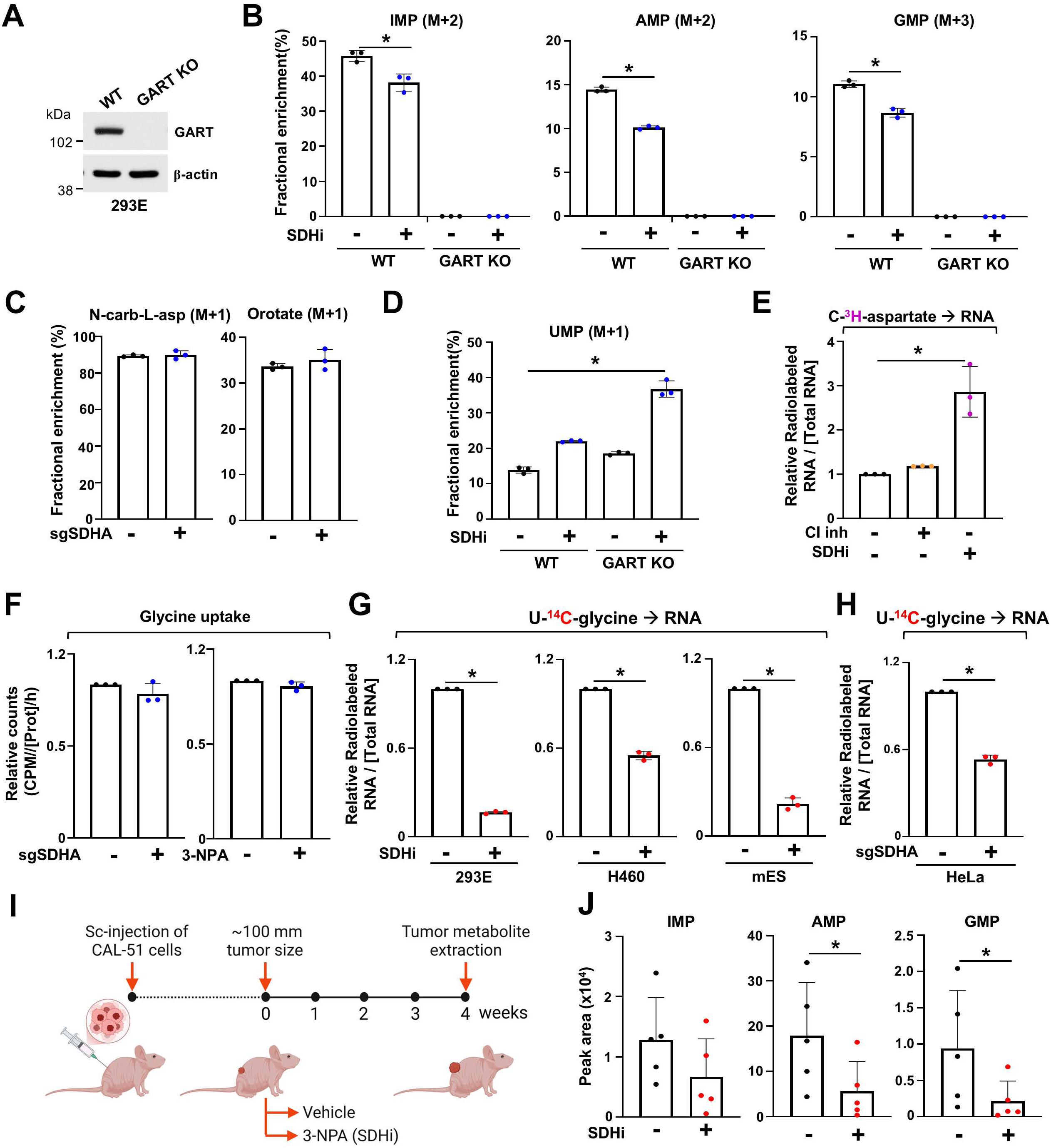
SDH inhibition leads to purine synthesis suppression. (A) Immunoblots of HEK 293E WT cells or knockout for the purine enzyme GART (*sgGART*). (B) Fractional enrichment (%) of the indicated metabolites from wildtype or GART KO HEK-293E cells and labeled with [^15^N-^13^C_2_]–glycine (400 µM) for 4 h and treated either with vehicle (DMSO) or 3- NPA. (C) Fractional enrichment (%) of the indicated pyrimidine intermediates from wildtype or SDHA KO (sgSDHA) HeLa cells labeled with [^15^N-(amide)]-glutamine (4 mM) for 2 hours. (D) Fractional enrichment (%) of UMP (M+1) from wildtype or GART KO HEK-293E cells treated with vehicle (DMSO) or 3-NPA (SDHi, 1 mM) and labeled with [^15^N-(amide)]-glutamine (4 mM) for 2 hours. (E) The relative levels of incorporation of [^3^H] from [C-2,3-^3^H]-aspartate into RNA are shown. Labeling in HeLa cells was performed for 6 hours treated with either vehicle (DMSO), rotenone (CI inh, 1µM), or 3-NPA (SDHi, 1 mM). Radioactivity measured reflects de novo pyrimidine synthesis activity. (F) Uptake of glycine in wildtype or SDHA KO HeLa cells or wildtype HeLa cells treated with vehicle (DMSO) or 3-NPA for 8 hours and labeled with [U-^14^C]-glycine for 5 min. (G) The relative levels of incorporation of [^14^C] from [U-^14^C]-glycine into RNA are shown. Labeling was performed in HEK 293E, H460 (lung cancer) and mouse embryonic stem (mES) cells, treated either with DMSO or 3-NPA for 9 hours. Radioactivity measured reflects de novo purine synthesis activity. (H) The relative levels of incorporation of [^14^C] from [U-^14^C]-glycine into RNA are shown. Labeling in wildtype and SDHA KO HeLa cells was performed for 4 hours. Radioactivity measured reflects de novo purine synthesis activity. (I) Schematic illustrating the generation of CAL-51 breast cancer subcutaneous xenografts in athymic nude mice and the treatment strategy. (J) Normalized peak areas of purine metabolites IMP, AMP, and GMP in vehicle (DMSO) and 3-NPA treated mice (n=5). For all plots, data are shown as mean ± s.d. of n= 3 independent biological replicates. \**P* < 0.05 were determined by unpaired, two-tailed Student’s t-test for pairwise comparisons (B, G-H, J) or one-way ANOVA with Tuckey post-hoc test for multiple comparisons (D, E).

**Supplementary Fig. 3.**
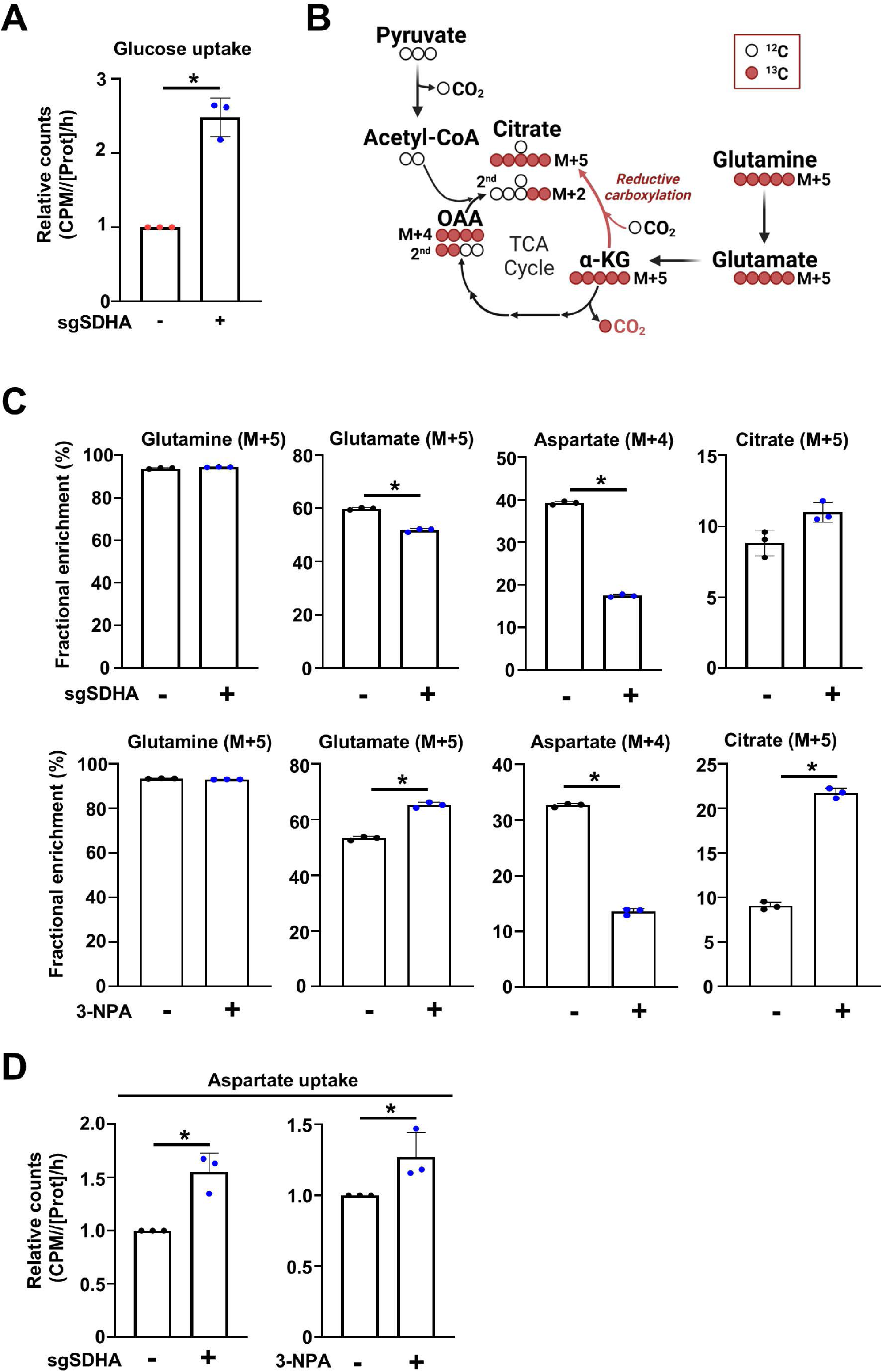
SDH inhibition stimulates glucose, aspartate uptake and glutamine-dependent reductive carboxylation. (A) Uptake of glucose measured in wildtype and SDHA KO HeLa cells cultured in 10% dialyzed serum. Cells were labeled with ^3^H-2-deoxyglucose over 5 min. The CPM values were normalized to the protein concentration and uptake duration. (B) Schematic illustrating the glutaminolysis pathway and reductive carboxylation. (C) Fractional enrichment (%) of glutamine (M+5), glutamate (M+5), aspartate (M+4), and citrate (M+5) from wildtype and SDHA KO HeLa cells or HeLa treated with either vehicle (DMSO) or 3-NPA (1 mM) cultured in 10% dialyzed serum and labeled with [^13^C_5_]–glutamine for the last 2 hours. (D) Uptake of aspartate measured in wildtype and SDHA KO HeLa cells or HeLa cells treated with vehicle or 3-NPA (1 mM) cultured in 10% dialyzed serum. Cells were labeled with ^3^H-aspartate over 5 min. The CPM values were normalized to the protein concentration and uptake duration. For all plots, data are shown as mean ± s.d. of n= 3 independent biological replicates. *P < 0.05 were determined by unpaired, two-tailed Student’s t-test (A, C-D).

**Supplementary Fig. 4.**
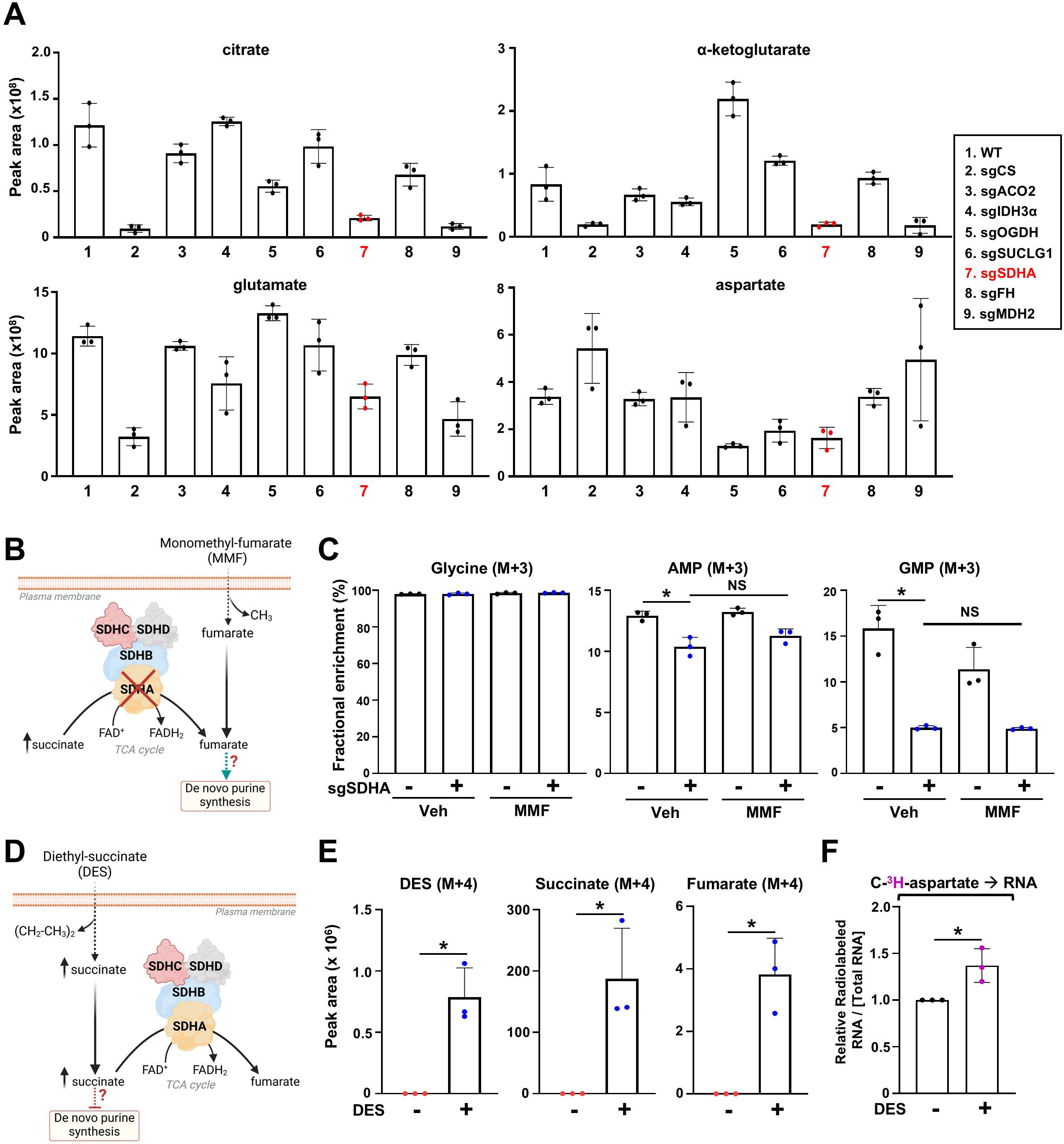
Fumarate supplementation fails to rescue de novo purine synthesis in SDH-deficient cells. (A) Normalized peak areas of citrate, a-ketoglutarate, glutamate, and aspartate in CS KO, ACO2 KO, IDH3α KO, OGDH KO, SUCLG1 KO, SDHA KO, FH KO and MDH2 KO HeLa cells cultured in 10% dialyzed serum. (B) Schematic demonstrating the entry of monomethylfumarate (MMF) and its conversion to fumarate in the cell. (C) Fractional enrichment (%) of glycine (M+3), AMP (M+3) and GMP (M+3) from HeLa cells treated either with vehicle (DMSO), MMF (5 mM) and labeled with [^15^N-^13^C_2_]–glycine for 4 h. (D) Schematic illustrating the entry of cell permeable succinate (diethyl-succinate (DES)) and its cellular conversion into succinate. (E) Normalized peak areas of DES (M+4), and TCA cycle intermediates from cells treated with vehicle (DMSO) or ^13^C_4_-diethyl-succinate (1mM) for 4 hours. (f) The relative levels of incorporation of [^3^H] from [2,3-^3^H]-aspartate into RNA are shown. Labeling was performed in HeLa cells treated with either vehicle (DMSO) or DES (1 mM) for 4 hours. Radioactivity measured reflects de novo pyrimidine synthesis activity. For all plots, data are shown as mean ± s.d. of n = 3 independent biological replicates. *P < 0.05 were determined by one-way ANOVA with Tuckey post-hoc test for multiple comparisons (C), or unpaired, two-tailed Student’s t-test for pairwise comparisons (E, F).

**Supplementary Fig. 5.**
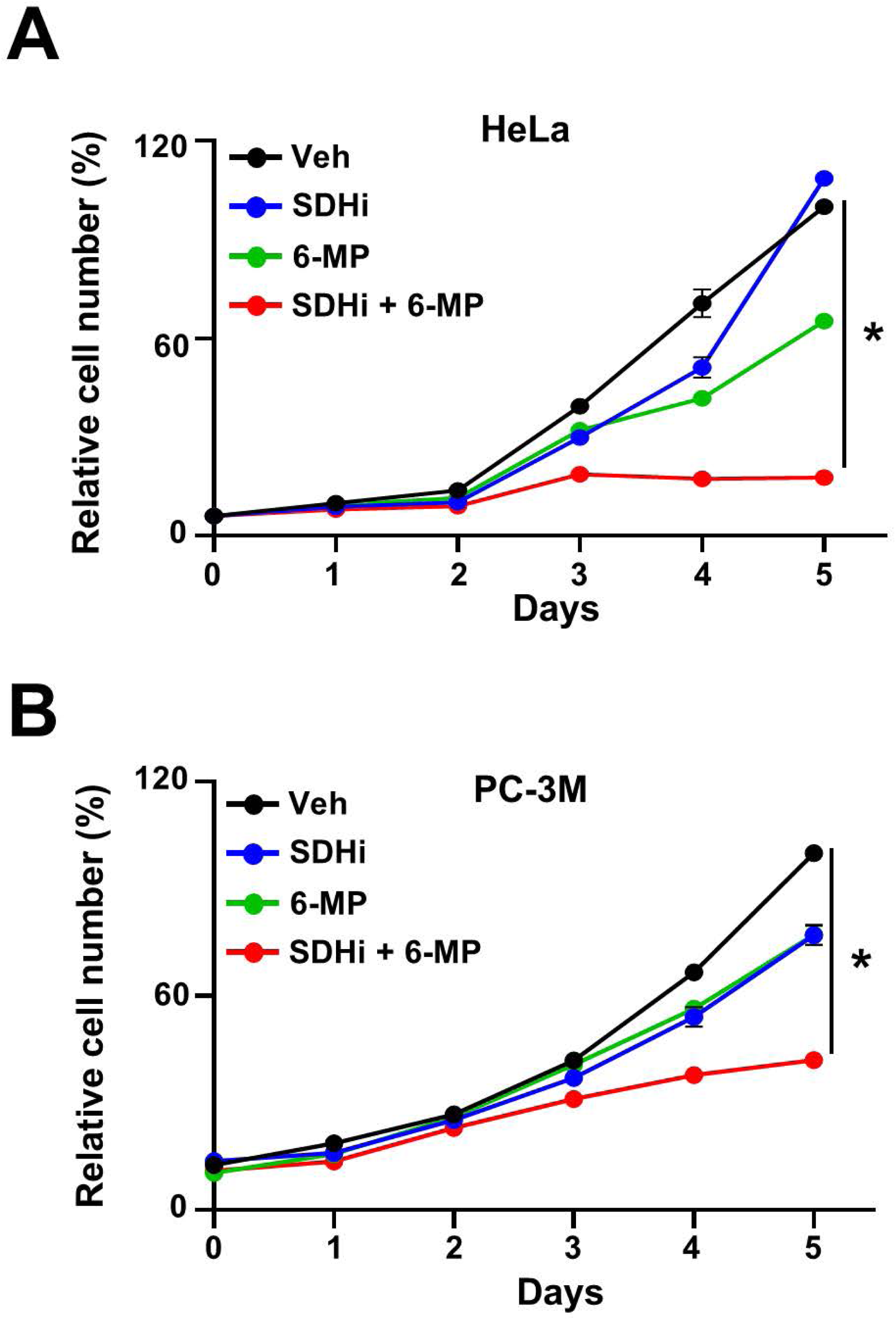
Co-targeting SDH and the purine salvage pathway decreases cancer cell growth. (A) HeLa cells treated with either vehicle, 3-NPA (100 µM), 6-MP (3 µM), or the combination, and crystal violet assays were performed every 24 h. (B) PC-3M cells treated with either vehicle, 3-NPA (30 µM), 6-MP (12.5 µM), or the combination, and crystal violet assays were performed every 24 h. * P < 0.05, by one-way ANOVA with Tukey’s post-hoc test (A, B).

## Acknowledgements

This research was supported by grants from the National Institutes of Health (NIH): R01GM135587, R01GM143334 (I.B.-S.), R01GM144617, R01CA258833, R21ES035975 (M.L.M.), R35GM137836 (N.S.); the LAM Foundation Established Investigator Award (LAM0151E01-22) (I.B.-S.); and American Cancer Society Awards: DBG-23-1039959-01-TBE (I.B.-S.), ABOA Impact RSG-22-086-01-TBE (M.L.M.). N.S. is a CPRIT Scholar in Cancer Research with funding from the Cancer Prevention and Research Institute of Texas (CPRIT) New Investigator Grant RR160021. N.S. was supported by the Andrew Sabin Family Foundation Fellowship. We thank Ram Prosad Chakrabarty and Navdeep Chandel for providing mouse embryonic stem cells. We are grateful to members of the Mendillo and Ben-Sahra laboratories at Northwestern University for their feedback on this study.

## Author contributions

I.B.-S. and M.L.M. conceived and supervised the study and drafted the manuscript. M.A.N. performed the experiments, analyzed the data, and contributed significantly to manuscript preparation. A.T.K. conducted the CRISPR screen analysis and cell biology experiments. H.S.C. U.S, and O.V.-C. assisted with additional cell biology and immunoblot assays. D.M. and N.S. engineered and established the CRISPR metabolic library. P.G., J.M.A., and H.S. carried out the LC-MS/MS analyses and provided expertise in mass spectrometry methodologies. M.A.N., I.B.- S., and M.L.M. collaboratively finalized the manuscript.

## EXPERIMENTAL MODEL AND SUBJECT DETAILS

Details of cell lines used have been provided under the section “Cell Lines and Tissue Culture” and details of mice used have been provided under “Allograft and xenograft experiments”.

### Cell Lines and Tissue Culture

HeLa, H460, CT26, 786-O, PC-3M, hTERT-RPE1, and HEK 293T cell lines were obtained from the American Type Culture Collection (ATCC), and CAL-51 cells were obtained from DSMZ. HEK293E cells were kindly provided by Dr. John Blenis (Weill Cornell Medicine). HeLa, HEK 293E, HEK 293T, CT26, and H460 cell lines were cultured in DMEM with 25 mM glucose (Fischer # 10-017-CV), 10 % Fetal bovine serum (Gibco # 26140-079), 2 mM glutamine (sigma # 59202C) 37°C, and 5 % CO_2_. CAL-51 cells were cultured in an analogous medium but with 20 % FBS. 786- O cells were cultured in DMEM high glucose supplemented with 5 mM sodium pyruvate (sigma # P2256-5G). hTERT-RPE1 cells were cultured in DMEM/F12 (Fischer # 11320033) supplemented with 10% FBS and 2 mM glutamine. PC-3M cells were cultured in RPMI-1640 supplemented with 10% FBS and 2 mM glutamine. Mouse embryonic stem cells were cultured in Serum-free 2i medium containing NEUROBASAL (Gibco # 21103-049), DMEM/F12 (Gibco # 11320-033), N2- SUPPLEMENT (Gibco # 17502-048), B27+RA (Gibco # 17504-044), 7.5% BSA (Gibco # 15260- 037) and PEN/STREP (corning # 30-001-CI). Media were changed to 10% dialyzed serum 15 hours before experiments. Viable cells were counted using a TC-20 Automated Cell Counter (Bio-Rad).

### DepMap CERES Score and FIREWORKS Coessentiality Network Generation

Using publicly available data from DepMap (24Q4), the CERES scores for all cancer cell lines for *SDHA*, *SDHB*, *SDHC*, *SDHD*, *GART*, *UMPS*, and *NDUFS1* were accessed and plotted^32^. A CERES score of 0 means that loss of a given gene has no fitness effect for a cell line, a positive CERES score indicates that loss of a given gene has a positive fitness effect, and a negative CERES score indicates that a gene has a negative fitness effect on a cell line, with a score of −1 corresponding to the median effect observed upon loss of a common essential gene.

Coessentiality network analysis was performed using FIREWORKS, a bias-corrected, rank-based coessentiality method and online tool^14^. Briefly, whole-genome CRISPR screening data from the Cancer Dependency Map (DepMap) was used to calculate the correlation of the CERES essentiality score between every pair of genes in the genome^32^. These correlations were processed with a bias-correction approach and can be used to construct rank-based coessentiality networks for any given gene (or combination of genes) as the source node(s). A solid red line represents a primary node that is positively coessential, and a dotted red line represents a secondary node that is positively coessential with a primary node. For SDHA, a coessentiality network of 20 positively correlated primary nodes with 5 positively correlated secondary nodes was generated.

### Constructs and CRISPR/Cas9

Full-length human SDHA cDNA in pCMV6-entry vector containing the FLAG-tag was purchased from Origene (RC200349). The SDHA sequence was cloned into a lentiviral compatible vector (MTHFD1 (NM_005956) Human Tagged Lenti ORF Clone, Origene # RC200297L3) using SgfI-MluI restriction digestion and cloning sites. To generate CS, ACO2, IDH3a, OGDH, SUCLG1, SDHA, FH, MDH2, and GOT2 knockout cells, single guide RNA (sgRNA) sequence targeting the specific genes were cloned into the lentiCRISPR v2 (Addgene, Plasmid # 52961) vector using the following oligonucleotides: CS 5’ AACTGGACATATCCCAACAG 3’, ACO2 5’ AGCGAGGCAAGTCGTACCTG 3’, IDH3α 5’ TAGCCTTCGATAGAGACACA 3’, OGDH 5’ GTTGGCCACTCATAGATACG 3’, SUCLG1: 5’ TCAGTGATACACACAACCAA 3’, SDHA: 5’ ACCGTGCATTATAACATGGG 3’, FH: 5’ CCAGTCTGCCATACCACGAG 3’, MDH2: 5’ CTGCCTGAAAGGTTGTGATG 3’, GOT2: 5’ AACAGCGAAGTCTTGAAGAG 3’.

### Lentivirus Production and Transduction

To generate SDHA cDNA virus, HEK293T cells were transfected with SDHA cDNA containing lentiviral vector along with the packaging and envelope plasmids (psPAX2 Addgene # 12260, pMD2.G Addgene # 12259) using lipofectamine 3000 transfection reagent (Fischer # L3000-015) according to the manufacturer’s guidelines. Similarly, HEK293T cells were transfected with lentiCRISPR v2 plasmid (Addgene # 52961), or with lentiCRISPR v2 plasmid containing sgRNA for target genes along with the packaging and envelope plasmids (psPAX2, pMD2.G) using lipofectamine 3000 transfection reagent according to the manufacturer’s guidelines. 8 hours after transfection, media were changed and 24 hours later, virus-containing media were collected and filtered at 0.45 μm and then immediately used to transduce cells. Cells were transduced with lentivirus containing 8 μg/mL polybrene (sigma # H9268). Sixteen hours after transduction, the media containing lentivirus was replaced with fresh media. Twenty-four hours later, the media was replaced with fresh media containing 2 μg/mL puromycin (Gibco # A1113803) for a total of 3 media changes.

### Immunoblotting

Cells were lysed in ice-cold Triton lysis buffer (120 mM NaCl, 40 mM HEPES, pH 7.4, 1% Triton X-100, 1 mM EDTA, 10 mM sodium pyrophosphate, 10 mM glycerol 2-phosphate, 0.5 mM sodium orthovanadate and 50 mM NaF; 1 μM Microcystin-LR and protease inhibitor cocktail were added) and incubated on ice for 30 min. Lysates were centrifuged at 20,000 g for 15 min at 4°C. Protein concentrations were quantified using Bradford assay and protein lysates were normalized. Equal amounts of protein lysates (∼10 to 20 μg) were prepared with the Laemmli sample buffer, and incubated at 95°C on a heat block for 5 min. Denaturated protein lysates were then separated on SDS-PAGE and transferred onto nitrocellulose blotting membranes. Membranes were blocked in 5 % non-fat milk in TBS tween (50 mM Tris pH 7.4, 150 mM NaCl, 0.1 % Tween 20) and incubated for 15 hours at 4°C with the indicated primary antibodies. The primary antibodies used target CS (Proteintech # 16131-1-AP), ACO2 (Proteintech # 11134-1-AP), IDH3α (Proteintech # 15909-1-AP), OGDH (Proteintech # 15212-1-AP), SUCLG1 (Proteintech # 14923-1-AP), SDHA (Proteintech # 14865-1-AP), FH (Proteintech # 11375-1-AP), MDH2 (Proteintech # 15462-1-AP), GOT2 (Proteintech # 14800-1-AP), GART (Proteintech # 13659–1-AP), β-actin (sigma # A5316), and FLAG M2 (sigma # F1804). Membranes were washed with TBST and incubated with horseradish peroxidase (HRP)-tagged anti-rabbit (CST # 7074S) or anti-mouse (CST # 7076S) secondary antibodies for one hour. Protein bands were developed with chemiluminescent substrates (Thermo Fisher Scientific # 34577, #34095). In all western blot experiments, β-actin was used as a loading control. All conditions were analyzed with biological triplicates and representative of at least three independent experiments.

### Cell proliferation assays

Cells were seeded in 96-well plates at a density of 1000-1500 cells per well in media containing 10 % dialyzed FBS. The next day, cells were treated as indicated in the figures. Cell proliferation was measured after regular intervals as indicated in the figures using 50 μL of 0.5% crystal violet staining solution (Fischer # AC405830250) to each well, and incubating for 20 min at room temperature on a bench rocker with a frequency of 20 oscillations per minute. The plates were washed four times in a stream of tap water and air-dried at room temperature. 200 μL of methanol was added to each well, and the plate was incubated with its lid on for 20 min at room temperature on a bench rocker with a frequency of 20 oscillations per minute. The optical density of each well was measured at 570 nm (OD_570_) with a plate reader. The data were normalized to vehicle control treated conditions. All cell proliferation assays were performed in biological triplicates seeded in technical quintuplicates for each condition.

### Compact Metabolic CRISPR Screen

We generated a library of sgRNAs targeting 232 metabolic genes (4 sgRNAs/gene) using the lentiGuide-Puro plasmid backbone (Addgene #52963). This library was transformed into Stbl3 bacteria and amplified at 30°C overnight. The library was then transfected into HEK-293T cells along with packaging plasmids pMD2.G (Addgene #12259) and psPAX2 (Addgene #12260) to generate lentivirus harvested from the media 48h and 72h after transfection.

The cell line used for the screen was immortalized retinal pigmented epithelial (hTERT- RPE1) cells grown in DMEM/F12 media supplemented with 10% heat-inactivated FBS and 1% Pen/Strep. Since the plasmid backbone for the library does not contain Cas9, RPE1 cells were transduced with lentivirus containing lentiCas9-blast (Addgene #52692) and selected with blasticidin at 10 μg/mL for 7 days to generate RPE1-cas9 cell line which has constitutive Cas9 expression. To determine the multiplicity of infection (MOI) of the metabolic library lentivirus, 3x 10^6^ RPE-cas9 cells in 15 cm^2^ dishes were transduced with a range of volumes of lentivirus. Media was changed one day after transduction and selection with 20 μg/mL of puromycin began the following day. After four days in puromycin, the cells from each plate were counted and compared to a no-virus kill control and a non-selected positive control. From this data, the dose that was closest to an MOI of 0.3 was calculated and was used for the screen.

For 3-NPA treatment, the IC_20_ was determined by seeding 5,000 RPE1-cas9 cells/well in a 96wp. The following day, a range of doses of the 3-NPA were applied. Three days later, 50 μL of 0.5% crystal violet staining solution (Fischer # AC405830250) was added to each well, followed by washing and air-drying the plates. 200 μL of methanol was added to each well and the optical density of each well was measured at 570 nm (OD570) with a plate reader. The data were normalized to vehicle control treated conditions.

For the screen, 12 x 15 cm^2^ dishes of RPE1-Cas9 were transduced at an MOI of 0.3 with the metabolic library lentivirus. Media was changed the following day with 20 μg/mL of puromycin added the day after. All plates were trypsinized and replated 48 hours after selection began to encourage further selection. Two days later (deemed T0), the selection was complete, and the remaining cells were expanded for 6 days until enough cells were generated to seed 3 x 10^6^ cells per plate and treatment condition. A pellet of 20 x 10^6^ cells was also snap-frozen at T0 for downstream analysis. The cells were then passaged every 3 days and fresh 3-NPA was added every passage. Cells were passaged until T21 at which point all the cells for a given condition were snap-frozen and stored at −80C. Cell pellets were thawed on ice for 15 minutes before genomic DNA extraction using the Puregene Cell Kit (Qiagen #158043) and following manufacturer instructions. Genomic DNA was then quantified by Qubit dsDNA BR (Invitrogen # Q32850). 20 μg of genomic DNA was used from each sample as an input for the PCR reaction to amplify the sgRNAs from the library. For each PCR reaction, 10 μL of 10 μM i5 and i7 primers, which target the U6 promoter of lentiGuide-Puro, were added to the genomic DNA with 100 μL of Q5 High-Fidelity 2X Master Mix (NEB # M0492S). The final reaction volume was brought up to 200 μL with water if necessary and each reaction was split into 4 x 50 μL reactions to avoid the possibility of PCR jackpot effects. Each reaction was then placed into a thermocycler (BioRad T100) with the following program: 98°C for 3 min, then 23 cycles of 98°C for 30 sec, 65.2°C for 30 sec, and a final extension of 72°C for 1 min with a 4°C hold. The 4 x 50 μL reactions were then re-pooled for each condition and run on a 1.5% agarose gel. A 363 bp band corresponding to the expected size of an amplified sgRNA was excised and purified by gel extraction following manufacturer instructions (Qiagen #28704). The purified library was then sequenced using a NovaSeq 6000 (Illumina) where each sample received a minimum of 10 x 10^6^ reads to maintain library coverage.

The reads from each sample were processed to count the number of sgRNAs present from the compact metabolic library using Model-based Analysis of Genome-wide CRISPR/Cas9 Knockout (MAGeCK)^33^. Once a count table was generated for all samples, drugZ, an algorithm for identifying both synergistic and suppressor chemogenetic interactions from CRISPR screens, was used to compare the 3-NPA-treated samples to the DMSO control and T0 conditions^34^. A normZ score (CRISPR viability score) was generated for each guide in the library with a negative normZ score corresponding to a depletion of a sgRNA in 3-NPA-treated cells Vs. vehicle control cells and a positive normZ score corresponding to an enrichment of a sgRNA. A false discovery rate (FDR) was also calculated for each sgRNA and an FDR < 0.2 was used as the cutoff for a significant hit from the screen.

### LC-MS based stable isotope tracing experiments

To determine the relative abundances of intracellular metabolites, extracts were prepared and analyzed via LC-MS/MS. Briefly, for targeted steady-state samples, metabolites were extracted on dry ice with 4 mL 80% methanol (−80°C), as described previously ^35^. Insoluble material was pelleted by centrifugation at 3000 g for 5 min, followed by two consecutive extractions of the insoluble pellet with 0.5 mL 80 % methanol, centrifugation at 20,000 g for 5 min. The 5-mL metabolite extract from the pooled supernatants was dried down under nitrogen gas using the N-EVAP (Organomation, Inc., Associates). 50% acetonitrile was added to the samples for reconstitution followed by vortexing for 30 sec. Samples solution was centrifuged 20,000 g for 30 min at 4 °C. Supernatant was collected for LC-MS analysis.

For isotope tracing experiments, cells were seeded in biological triplicate (∼70–80% confluent), washed once with serum-free DMEM, and then incubated with either 10 mM [^13^C_6_]- glucose (Cambridge Isotope Laboratories # CLM-1396-2), 4 mM [^15^N-(amide)]-glutamine (Cambridge Isotope Laboratories # NLM-557-1), 4 mM [^13^C_5_]-glutamine (sigma # 605166-100MG) or 400 μM [^15^N-^13^C_2_]-glycine (Cambridge Isotope Laboratories # CNLM-1673-H-0.5) in either glucose-free, glutamine-free, or glycine-free media, respectively. Metabolites were extracted as described in the steady-state studies. For both steady-state and isotope tracing experiments, samples were analyzed by High-Performance Liquid Chromatography, High-Resolution Mass Spectrometry, and Tandem Mass Spectrometry (HPLC-MS/MS). In detail, the LC-MS/MS system is comprised of a Thermo Q-Exactive in line with an electrospray ionization (ESI) source and an Ultimate3000 (Thermo) series HPLC consisting of a binary pump, degasser, and auto-sampler outfitted with an Xbridge Amide column (Waters; dimensions of 2.3 mm × 100 mm and a 3.5 μm particle size). The mobile phase A contained 95% (vol/vol) water, 5% (vol/vol) acetonitrile, 10 mM ammonium hydroxide, 10 mM ammonium acetate, pH = 9.0; B was 100% Acetonitrile. The gradient was as follows: 0 min, 15 % A; 2.5 min, 30 % A; 7 min, 43 % A; 16 min, 62 % A; 16.1– 18 min, 75 % A; 18–25 min, 15 % A with a flow rate of 150 μL/min. The capillary of the ESI source was arranged for 275 °C, with sheath gas at 35 arbitrary units, auxiliary gas at 5 arbitrary units, and the spray voltage at 4.0 kV. In positive/negative polarity switching mode, an m/z scan range from 60 to 900 was chosen and MS1 data was collected at a resolution of 70,000. The automatic gain control (AGC) target was set at 1 × 10^6^ and the maximum injection time was 200 ms. The top 5 precursor ions were then fragmented, in a data-dependent manner, using the higher energy collisional dissociation (HCD) cell set to 30% normalized collision energy in MS2 at a resolution power of 17,500. Besides matching m/z, metabolites are identified by matching either retention time with analytical standards and/or MS2 fragmentation pattern. Data acquisition and analysis were carried out by Xcalibur 4.1 software and Tracefinder 4.1 software, respectively (both from Thermo Fisher Scientific). All conditions were analyzed with biological triplicates and representative of at least three independent experiments.

### Measurement of [^14^C]-glycine, [C^3^H]-aspartate, and [^3^H]-hypoxanthine incorporation into nucleic acids

75-80% confluent cells were cultured in 10% dialyzed serum for 15 hours, then treated and labeled as indicated in the figures. Cells were labeled with 2 μCi of [U-^14^C]-glycine (ARC #ARC 0292-250 µCi), [C^3^H]-aspartic acid (ARC # ART 0211-250 µCi), [^3^H]-Hypoxanthine Monohydrochloride (Perkin Elmer # NET177001MC). RNA or DNA was purified using an Allprep DNA/RNA mini kit (Qiagen # 80204) according to the manufacturer’s instructions and quantified using a spectrophotometer. 30 μl of eluted RNA were added to scintillation vials containing 3 mL of liquid scintillation emulsifier safe (Perkin Elmer LLC Emulsifier Safe Cat N°50-905-0578) and radioactivity was measured by liquid scintillation counting and normalized to the total RNA or DNA concentrations, respectively. All conditions were analyzed with biological triplicates and representative of at least three independent experiments.

### Glucose, glycine, and aspartate uptake measurements

Uptake was measured using a modification of the amino acid uptake protocol as previously described (Edinger and Thompson, 2002). Briefly, HeLa cells were cultured in serum-free DMEM for 15 hours and 50 μL of uptake medium containing 1 μCi of specific activity for either 2-[1,2-^3^H]-deoxyglucose (ARC #ART 0103-250 µCi), U-^14^C-glycine (ARC #ARC 0292-250 µCi), or L-[2,3-^3^H]-aspartic acid (ARC # ART 0211-250 µCi), was added for 5 min at 37°C. The uptake was stopped promptly by washing the cells in ice-cold PBS (Fischer # 14190250) and lysing them in Triton 1 % buffer (40 mM HEPES, pH 7.4, 120 mM NaCl, 1 mM EDTA, 1 % Triton X-100, 10 mM glycerol 2-phosphate, 10 mM sodium pyrophosphate, 0.5 mM sodium orthovanadate and 50 mM NaF; protease inhibitor cocktail was added before the lysis of cells). Samples were centrifuged at 20,000 g and 4°C for 15 min. 75% of the supernatant was added to scintillation vials and radioactivity was measured by liquid scintillation counting (c.p.m) and normalized to the protein concentrations. Data points are presented as the means of triplicate samples ± s.d.

### Mouse allograft and xenograft experiments

All animal procedures and studies were approved by the Institutional Animal Care and Use Committee (IACUC) at Northwestern University. All experiments were performed following relevant guidelines and regulations. 5-week-old BALB/c mice (Female) were injected subcutaneously in the flanks with the murine cancer cell line CT-26 (2 × 10^6^ cells) in a 1:1 mixture of Cultrex Basement Membrane Extract, Type 3, Pathclear (R&D Systems # 3632-001-02) and 0.9% saline. 5-week-old Athymic nude mice (Female) were injected subcutaneously in the flanks with th human cancer cell line CAL-51 cells (2 × 10^6^ cells) in a 1:1 mixture of Cultrex Basement Membrane Extract, Type 3, Pathclear (R&D Systems # 3632-001-02) and 0.9% saline. When tumors reached a size of approximately 100 mm^3^, the mice were treated with intraperitoneal injections for five days a week. Each mouse received no more than 0.5 μL/g DMSO per dosing. Tumor size was measured in three dimensions using an electronic caliper twice per week for four weeks and mice’s body weight was recorded every week. Mice were fasted overnight and injected with 3-NPA (30 mg/kg) in the morning 2 hours before in vivo stable isotope tracing. In vivo, stable isotope tracing experiments were performed by intraperitoneal bolus injections of [^15^N-^13^C_2_]- glycine (sigma # 489522) (0.5 g/kg) dissolved in 0.9% saline. Mice were euthanized 4 hours after the last injection. The tumors were harvested, frozen in ice-cold methanol 80 %, and stored at −80°C until samples were ready for processing.

### Quantification and statistical analysis

One-way ANOVA followed by Tukey’s post hoc tests was performed in GraphPad Prism 9.0 to determine differences between each group when more than two conditions were present. A two-tailed Student’s t-test was performed in GraphPad Prism 9.0 for two pairwise comparisons. All error bars represent the standard error of the mean (s.d.). A value of p < 0.05 was considered significant. The data represented is best from more than two independent experiments performed.

